# Modelling dynamic host-pathway interactions at the genome scale with machine learning

**DOI:** 10.1101/2024.04.09.588720

**Authors:** Charlotte Merzbacher, Oisin Mac Aodha, Diego A. Oyarzún

## Abstract

Pathway engineering offers a promising avenue for sustainable chemical production. The design of efficient production systems requires understanding complex host-pathway interactions that shape the metabolic phenotype. While genome-scale metabolic models are widespread tools for studying static host-pathway interactions, it remains a challenge to predict dynamic effects such as metabolite accumulation or enzyme overexpression during the course of fermentation. Here, we propose a novel strategy to integrate kinetic pathway models with genome-scale metabolic models of the production host. Our method enables the simulation of the local nonlinear dynamics of pathway enzymes and metabolites, informed by the global metabolic state of the host as predicted by Flux Balance Analysis (FBA). To reduce computational costs, we make extensive use of surrogate machine learning models to replace FBA calculations, achieving simulation speed-ups of at least two orders of magnitude. Through case studies on two production pathways in *Escherichia coli*, we demonstrate the consistency of our simulations and the ability to predict metabolite dynamics under genetic perturbations and various carbon sources. We showcase the utility of our method for screening dynamic control circuits through large-scale parameter sampling and mixed-integer optimization. Our work links together genome-scale and kinetic models into a comprehensive framework for computational strain design.

## I. INTRODUCTION

Microbial cell factories use the native metabolic capabilities of microorganisms to produce high-value compounds for a range of sectors such as energy, pharmaceuticals, and agriculture^1–3^. These production systems are built by expressing enzymatic genes in a suitable cellular host that catalyzes a conversion route from native precursors to a target compound. However, heterologous pathways interact with native cellular processes in a myriad of ways that impact the metabolic capacity and viability of the host. These effects occur across multiple cellular systems and scales, e.g. by consuming energy resources and cofactors, producing toxic intermediates, or directing metabolic flux away from bottlenecks in central metabolism^4^. As a result, pathway design requires a careful balance of enzyme expression levels, production yield and the resulting metabolic burden on the host.

Pathway design typically relies on genome-scale metabolic models (GEMs) to tease out host-pathway interactions and predict their impact on production. To date, GEMs have been built for thousands of organisms, including many bacterial species commonly employed as production hosts^5,6^. The host GEM is typically augmented with a target pathway^7–9^ that can be analyzed with many available software tools^10^. Among these techniques, Flux Balance Analysis (FBA) is a particularly well-adopted framework to predict genome-scale flux distributions under the assumption of a steady state optimality principle, which is typically taken to be growth rate, production yield, or a combination of both^7^.

A key gap, however, is the lack of tools for predicting the dynamics of host-pathway interactions, i.e. those effects that depend on the temporal dynamics of pathway enzymes and metabolites during the course of fermentation. For example, pathway bottlenecks can form during culture and toxicity effects can be triggered by the transient accumulation of pathway intermediates. Other examples include the use of intracellular control systems to dynamically adjust pathway expression levels^11^, or the use of external inducers to switch across pathways during fermentation^12^. In the synthetic biology literature, dynamic models for host-circuit interactions have been subject of much attention, including several approaches to model competition for shared cellular resources^13–16^ or the impact of growth feedback on circuit function^17^. Yet so far these approaches are limited to predictions on gene expression alone and do not include metabolic dynamics.

Predicting the dynamics of host-pathway interactions requires multi-scale models that simultaneously describe local dynamics of pathway metabolites and cofactors, heterologous enzymes, and their impact on the global metabolism of the expression host. Traditional FBA cannot account for metabolite dynamics because it assumes that metabolites are at steady state and, moreover, it works on the space of fluxes without information on the enzyme kinetics that support them. Early proof-of-concept studies attempted to predict native metabolite dynamics using small stoichiometric models of specific pathways such as central carbon metabolism^18–20^. Early works also included temporal dynamics of metabolic fluxes using FBA in combination with dynamic optimization algorithms^21^. Such dynamic FBA approaches model the dynamics of exchange fluxes and extracellular metabolites by combining a FBA predictions with a bioreactor model based on differential equations. Recent studies have proposed to integrate machine learning predictors into dynamic FBA algorithms for improved predictivity^22,23^. Other related studies have built variations of dynamic FBA to include dynamics of enzyme expression^24–27^, yet these approaches cannot predict intracellular metabolite concentrations or incorporate transcriptional regulatory loops commonly employed for dynamic pathway control^28^.

A complementary approach is kinetic modelling based on Ordinary Differential Equations (ODEs). Several works have expanded such models to the genome scale^29–33^ with some success in predicting dynamics in model microbes as well as detecting genetic interventions that improve yield^34^. Such models, however have hundreds of parameters that are challenging to fit to limited experimental data and often vary by multiple orders of magnitude across conditions and species^35^, which can limit their applicability^36^.

Here, we present a novel strategy for predicting pathway dynamics coupled with the metabolism of a production host. The approach produces local simulations of enzyme and metabolite concentrations informed by the global metabolic state of the host. We employ a combination of kinetic models and FBA simulations, whereby both models iteratively pass information to each other along the temporal trajectory. This requires computing FBA solutions at many points in time and thus leads to a prohibitively long simulation runtime. We resolved this challenge with surrogate machine learning models to replace most FBA calculations. By pre-training the surrogate models with offline FBA solutions, we were able to improve computational speed by at least 100-fold. We illustrate the effectiveness of our approach using two production pathways from the literature^37,38^ and the latest genome-scale model of *Escherichia coli* ^39^. We showcase the application of our method to predict metabolite dynamics in response to genome-wide knockouts and various carbon sources. We finally demonstrate the utility of the approach for the design of dynamic control circuits via large-scale sampling and global optimization. Our results provide a novel approach for computational strain design that includes temporal effects and the cross-talk between a pathway and the host where it resides.

## II. RESULTS

### A. Integration of genome-scale and kinetic models

We focus on the simulation of heterologous pathways that interact with the host metabolism through sharing of metabolites and/or co-factors. As illustrated in Figure 1A, our strategy relies on successive iterations between two models: a dynamic Ordinary Differential Equation (ODE) model for a target heterologous pathway, and a static, genome-scale, model for the host metabolism. The approach is centered on the ODE model to obtain local predictions of the temporal dynamics of pathway metabolites and enzymes, using the global metabolic state of the host as predicted by Flux Balance Analysis (FBA). Specifically, we first consider a kinetic model for the heterologous pathway:

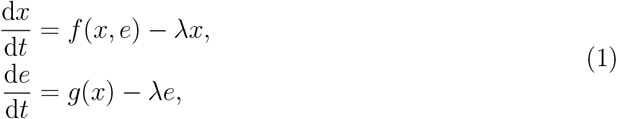

where *x* and *e* are vectors of metabolite and enzyme concentrations, respectively, and the rate constant *λ* models the dilution effect caused by cell growth. The vector function *f* (*x, e*) models the mass balance relationships and enzyme kinetics, and *g*(*x*) is a lumped model for the transcriptional and translational processes that control expression of pathway enzymes. This function can be used to flexibly accommodate for either constitutive enzyme expression, in which case *g*(*x*) is a constant, or metabolite-dependent enzyme expression as commonly encountered in dynamic pathway engineering^28^, in which case *g*(*x*) can be modelled as a Hill function of a pathway metabolite^40^.

**FIG. 1.**
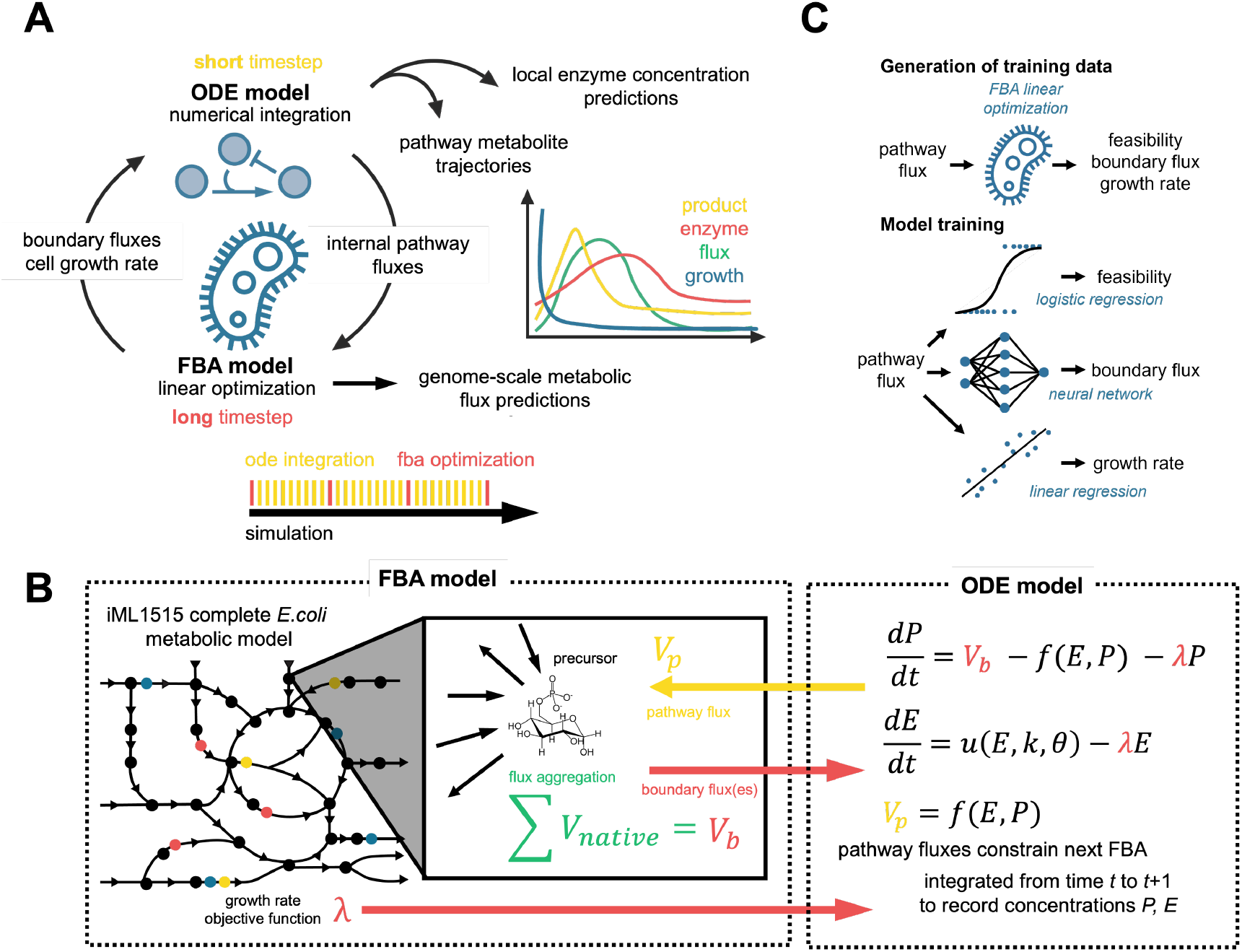
Schematic of our strategy to model dynamic host-pathway interactions. **(A)** The Flux Balance Analysis (FBA) model generates an optimal growth rate and boundary fluxes that feed into the Ordinary Differential Equation (ODE) model. The ODEs are then numerically integrated (short timestep) until the following long timestep, which then generates constraints on specific pathway consumption fluxes for FBA. Altogether, the FBA models generates growth rate and genome-scale flux predictions, while the ODEs generate local predictions for pathway metabolites and enzymes. Inset: FBA is run at every second of simulation (red) and the ODE solver integrates at many small, adaptively sized timesteps (yellow) in between these longer timesteps. **(B)** Detailed description of simulator. The iML1515 genome-scale model for *Escherichia coli* generates the growth rate and native metabolic fluxes. The native metabolic fluxes which generate or consume the pathway precursors are aggregated into a vector of boundary fluxes *V*_b_ to be passed to the ODE system. Upon integration, the ODE system generates a pathway flux, *V*_p_, which constrains further runs of the linear optimization in the FBA model. Colored dots in the metabolic network represent enzymatic cofactors that participate in many reactions. **(C)** Schematic of machine learning surrogates to replace the FBA calculations and increase simulation speed. We trained a logistic regression model to predict FBA feasibility. Feasible samples are then used to train a neural network to predict boundary fluxes and a linear regressor to predict growth rate.

The second model corresponds to a standard FBA formulation applied to a genome-scale model of the host:

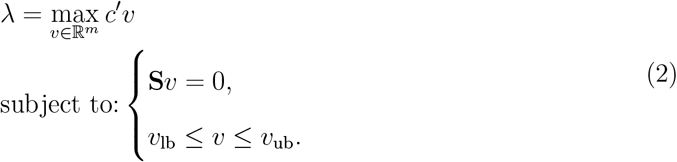

In this FBA problem, *λ* is the solution of a linear optimization problem with growth rate set as the objective function to be maximized^7^. In Eq. (2), **S** is the stoichiometric matrix of the host metabolism, *v* is a vector of *m* metabolic fluxes, *c* is a weight vector that describes the contribution of key reactions to biomass formation, and (*v*_lb_, *v*_ub_) are vectors of lower and upper bounds for each metabolic flux. The stoichiometric relation **S**_*v*_ = 0 imposes a steady-state constraint for all metabolites included in the genome-scale model.

The iterations between both models (Figure 1A) follow these principles: First, the FBA model in Eq. (2) produces a vector of fluxes *V*_b_ (which we call the “boundary fluxes”) and a predicted growth rate *λ*, all of which are passed as parameters to the ODE model in Eq. (1). Second, we integrate the ODE model on a short time interval to produce temporal predictions of metabolite and enzyme concentrations. Third, at the end of the integration interval, we compute a flux *V*_p_ consumed by the pathway from the host (which we call the “pathway flux”) that gets passed back to the FBA model as an equality constrain. By iterating these steps, the algorithm produces temporal trajectories for fluxes, metabolites, and enzymes of the heterologous pathway, alongside the dynamics of growth rate of the host. This approach to jointly simulate FBA-ODE models operates on two timescales: a fast metabolic timescale comprised in the FBA problem and kinetics of the production pathway, and a slow timescale of heterologous enzyme expression and cell growth.

In our formulation, the heterologous pathway is not included in the genome-scale model but instead simulated separately. We implemented this by adding a sink reaction with pathway flux *V*_p_ to the genome-scale model with stoichiometry *P* → ∅, where *P* is the host precursor that feeds into the heterologous pathway. We note that boundary fluxes present in both models must satisfy the steady-state constraint imposed by FBA at each iteration of the simulation loop, so the ODEs are modified to ensure this relationship holds. Another challenge is the selection of initial metabolite and enzyme concentrations for the ODE simulation, as these can produce a pathway flux *V*_p_ that violates the GEM constraints.

We resolved this through a warm-up routine which iteratively finds initial conditions using a Bayesian optimization method^41^. Details on the balancing equations and the warm-up routine can be found in the Methods.

### B. Surrogate models using machine learning

While the initial algorithm design produced promising results and converged to realistic phenotypes, we found that simulation run times were prohibitively long for practical use, (e.g. several hours of runtime to generate a single 24h pathway simulation). We performed a runtime analysis on a preliminary model and found that the FBA optimization step was responsible for more than 95% of runtime (Supplementary Table S4). To accelerate the simulation, we employed surrogate machine learning models to replace the FBA steps (Figure 1C). Specifically, we first trained a binary classification algorithm to determine whether a given pathway flux (*V*_p_) resulted in a feasible FBA problem. An FBA problem is infeasible when there is no flux vector that satisfies the GEM constraints. We trained a logistic regression classifier to predict a binary outcome (1 if feasible; 0 if infeasible) from *V*_p_ values, so as to perform the growth rate and pathway flux predictions only for feasible samples.

For samples classified as feasible, we trained a feedforward neural network to predict each component of the boundary flux vector *V*_b_, and a linear regressor to predict the optimal growth rate *λ*. All regressors were trained on data pairs (*V*_p_, *V*_b_) and (*V*_p_, *λ*), respectively, generated through a large collection of offline FBA simulations; these data need to be generated only once per genome-scale metabolic model, thus leading to substantial gains in performance as compared to online FBA simulations run at every iteration. The surrogate models accelerated our simulations by more than 100-fold (Supplementary Figure S2), which enabled prediction of long time courses. The surrogate models achieved highly accurate performance (over 98% feasibility classification accuracy, Supplementary Figure S3) even with relatively small training sets (*N* = 250). We employed the speed-up enabled by the surrogate models to explore the impact of various perturbations described next.

### C. Dynamic host-pathway simulations in *Escherichia coli*

We employed our method in two cases studies involving production of glucaric acid^42^ and beta-carotene^43^ in *Escherichia coli*, using its latest genome-scale metabolic model (iML1515)^39^. Both pathways are representative of real-world applications of our simulation approach, as they contain nonlinear enzyme kinetics and regulatory loops, and feed from precursors from central and secondary metabolism. Details on model construction and parameter values can be found in the Methods and Supplementary Information.

#### a. Synthesis of glucaric acid

We first focused on a pathway that branches off of central carbon metabolism, and hence is expected to interact strongly with the host metabolism. Glucaric acid is a key precursor for a number of biomedical applications and its pathway has been implemented previously in *E. coli* ^42^. As shown in Figure 2A, the pathway requires four reactions: the first step (inositol-3-phosphate synthetase, Ino1, taken from *S. cerevisiae*) catalyzes glucose-6-phosphate (G6P) from central carbon metabolism into myoinositol-1-phosphate. This reaction is the pathway flux *V*_p_ which constrains the FBA model. Myoinositol-1-phosphate is subsequently converted into glucaric acid by inositol-1-monophosphatase (SuhB, native to *E. coli*), myoinositol oxidase (MIOX, from *Mus musculus*) and uronate dehydrogenase (Udh, from *Pseudomonas syringae*). In addition to being exported from the cell and acting allosterically on MIOX activity, myoinositol can sequester the transcription factor IpsA, a dual transcriptional regulator from *Corynebacterium glutamicum*. This feature can be exploited to build dynamic control circuits that respond to the intracellular levels of myoinositol, for example by using IpsA to control expression of Ino1 or MIOX. For the representative simulation shown in Figure 2B, we use a dual control genetic feedback architecture where IpsA represses the upstream enzyme Ino1 and activates the downstream enzyme MIOX, creating two negative feedback loops^38^. The sole boundary flux *V*_in_ is a sum of all reaction fluxes that produce and consume G6P in the genome-scale model, and *V*_p_ is the flux of the reaction catalyzed by Ino1.

**FIG. 2.**
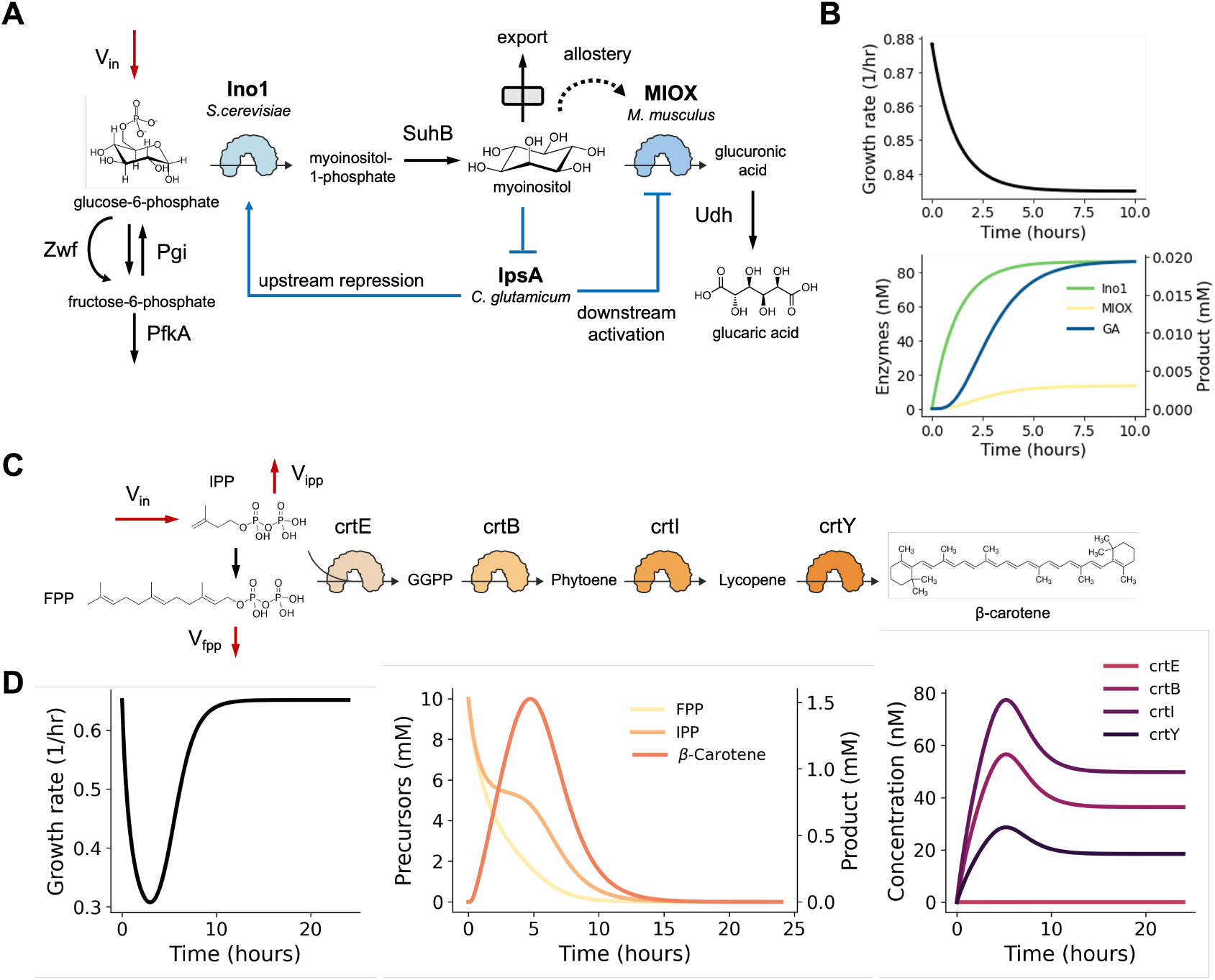
Pathway case studies in *Escherichia coli*. **(A)** Glucaric acid (GA) production pathway. Glucose-6-phosphate is converted to glucaric acid via myoinositol (MI). The enzyme MIOX is activated allosterically by its substrate MI and MI is also exported from the cell. The heterologous enzymes Ino1 and MIOX can be regulated via a metabolite-responsive transcription factor, IpsA, that can be repressed upon binding to MI. Regulatory architectures mediated by IpsA can repress Ino1 transcription upstream or activate MIOX transcription downstream, or both in a dual control architecture^44^ (shown in blue). The boundary flux *V*_in_ computed from FBA is labeled in red. **(B)** Simulation of host-pathway dynamics for glucaric acid production. The host-pathway model was simulated for 10 hours of simulation time (approx. 2.5 minutes of runtime) under a dual control architecture with parameters given in Supplementary Table S3. **(C)** Beta-carotene production pathway. Four heterologous enzymes (crtE, crtB, crtI, and crtY) produce beta-carotene from the precursors isopentyl pyrophosphate (IPP) and farnesyl pyrophosphate (FPP). The three boundary fluxes computed from FBA are labeled in red. **(D)** Simulation of host-pathway dynamics for beta-carotene production. The host-pathway model was simulated for 24 hours of simulation time (approx. 5 minutes of runtime) with parameters given in Supplementary Table S3. The beta-carotene pathway model was constructed from first principles using kinetic parameters sourced from literature or estimated with the deep learning model from Li *et al*. Details on model equations and parameters for both pathways are given in the Methods and Supplementary Information.

Glucose-6-phosphate is the product of the first step of glycolysis and catalyzed directly from glucose, the standard carbon source for *E. coli*. Genome-scale models for *E. coli* include G6P in the FBA objective function because of its direct impact on growth rate. We integrated a previously published kinetic model of the GA pathway^44^ with the iML1515 genome-scale model. Our host-pathway simulations indeed show a sustained drop in growth rate (Figure 2B), in line with the centrality of the precursor G6P. While the drop in growth rate depends on the expression levels of the heterologous enzymes, we generally observed realistic growth defects of 5–50% from the wild type^37,45^. We also observed a reasonable timescale of growth rate dynamics: a steady-state growth rate was achieved within 10 hours and 90% of the drop in growth rate occurs in the first 6 hours^46^.

#### b. Synthesis of beta-carotene

As a second example that branches from a down-stream region of metabolism, we selected the beta-carotene production pathway. We chose this pathway because it branches from native metabolites not included in the biomass equation and thus is not directly coupled to growth in the FBA formulation. Beta-carotene is a high-value metabolite with applications in medicine, cosmetics, animal feed, and nutritional supplements^47,48^. The biosynthetic beta-carotene production pathway has been implemented experimentally^49–51^ using codon-optimized enzymes from *Erwinia uredovora* engineered into *E. coli*. The pathway has four enzymes: crtE, crtB, crtI, and crtY (see Figure 2C). The two precursors farnesyl pyrophosphate (FPP) and isopentyl pyrophosphate (IPP) are catalyzed by crtE to form geranylgeranyl pyrophosphate (GGP). The enzyme crtB then catalyzes the conversion to phytoene, followed by crtI converting phytoene to lycopene. The enzyme crtY catalyzes the final step in the pathway to produce beta-carotene.

Unlike the glucaric acid pathway, which has only one boundary flux (*V*_in_ in Figure 2A), the beta-carotene model combines two precursors, FPP and IPP, both of which have their own effluxes and influxes. As shown in Figure 2C, the boundary flux vector *V*_b_ is thus composed of three elements: *V*_in_ is the influx to IPP, *V*_ipp_ is the efflux from IPP, and *V*_fpp_ is the efflux from FPP. The pathway flux *V*_p_ is the crtE-catalyzed conversion from the precursors to GGPP.

We built a kinetic model for the beta-carotene pathway using a combination of published parameter values and *k*_cat_ predictions from a deep learning model^52^, and integrated it with the iML1515 genome-scale model. Our host-pathway simulations (Figure 2D) show that in this case pathway activity leads to a transient growth defect, in line with the fact that precursors FPP and IPP are not weighed in the FBA objective function. Production of beta-carotene results in a defect to growth rate which recovers upon consumption of the precursors and reduction of pathway flux. The metabolite dynamics include consumption of the precursors followed by peaks of each intermediate in the pathway. The timing and size of these peaks depends on the enzyme promoter strengths, and we generally observed beta-carotene concentrations in a feasible mM range. The enzyme expression dynamics are also realistic, taking over 10 hours to reach steady state, and operate in a feasible nM range. Altogether, both case studies (glucaric acid and beta-carotene) demonstrate the reasonable results produced by our simulation approach and its applicability to a wide range of metabolic pathways.

### D. Growth and production in different carbon sources

Since *E. coli* can grow in a variety of carbon sources, we sought to predict the impact of growth media on the temporal dynamics of the glucaric acid pathway in Figure 2A. We chose a panel of 15 carbon sources that enter central metabolism at different points, both upstream and downstream of glucose-6-phosphate, which feeds into glucaric acid synthesis (Figure 3A). To perform these simulations, the machine learning FBA surrogates had to be re-trained for each of the 15 growth conditions, using data produced with offline FBA simulations. Details on data generation and model training can be found in the Methods.

**FIG. 3.**
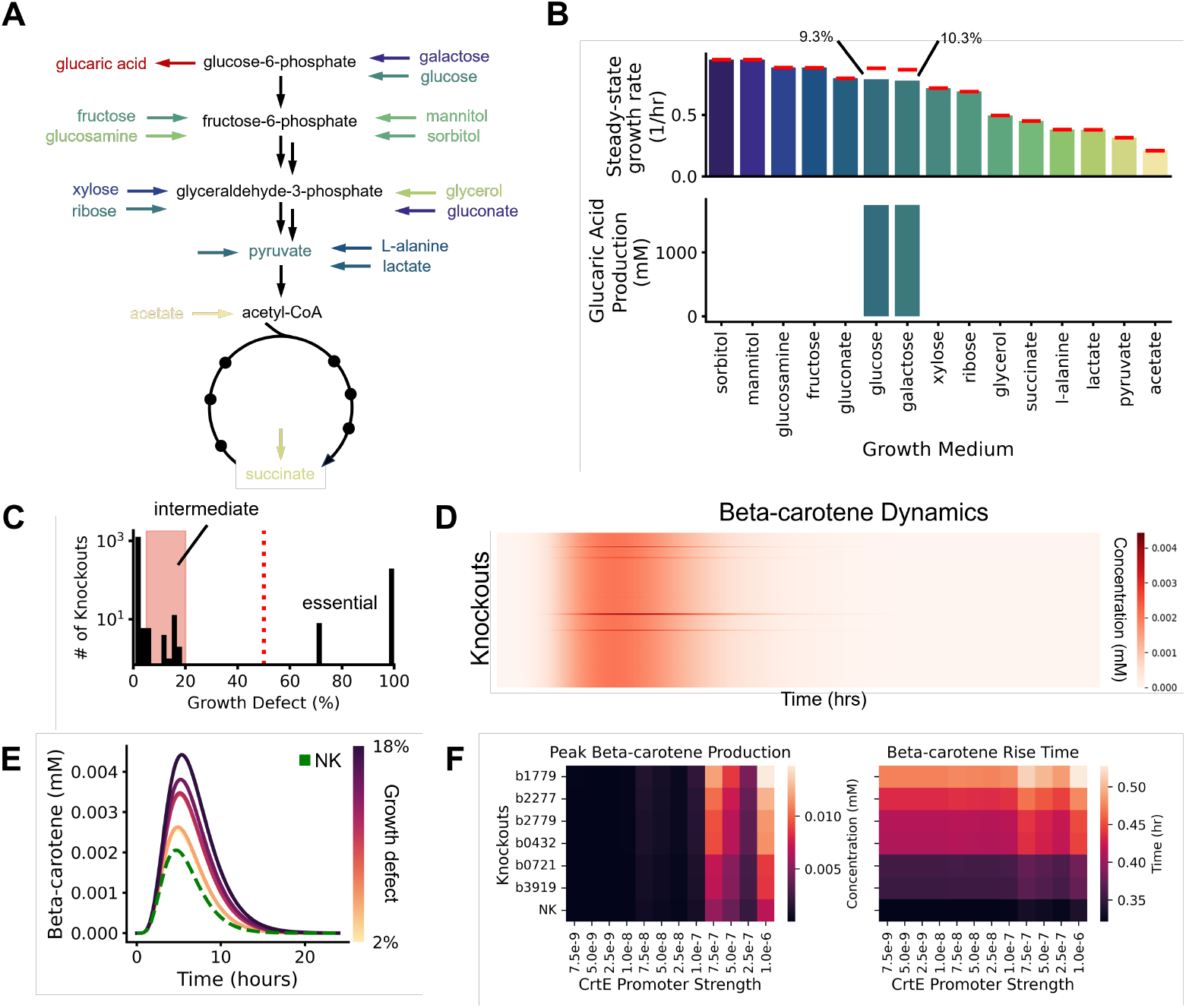
Consistency of simulation results under media and genetic perturbations. **(A)** Carbon sources entering at different points in central carbon metabolism (CCM). As shown in Figure 2A, the glucaric acid (GA) production pathway branches from CCM at glucose-6-phosphate (G6P). Besides galactose and glucose, all considered carbon sources enter CCM below G6P and therefore cannot be used for GA production. **(B)** Predicted steady-state growth rates and cumulative GA production, computed as the temporal integral of the GA production flux. Simulations were run for 24 hours of simulation time with parameters given in Supplementary Table S3. Wild type growth rates predicted by FBA are shown as red lines; our simulations predict GA production and a resulting growth defect in glucose and galactose. **(C)** Genome-scale knockout screen of strains engineered with the beta-carotene pathway in Figure 2C. The bar plots show a histogram of all knockouts and their resulting growth defect. The majority of knockouts either have no effect on growth, i.e. 0% growth defect from wild type (0.651/h), or are essential, i.e. >50% defect shown by red dotted line. Knockouts with a defect in the range 5–20% are defined as intermediate knockouts and highlighted in red. Simulations were run for 24 hours of simulation time with parameters given in Supplementary Table S3. **(D)** Beta-carotene concentration dynamics for all nonessential knockouts. Each row of the heatmap is a timecourse of beta-carotene production. Darker colors indicate a peak in the production. Knockouts with significant impact on product dynamics can be seen as streaks on the plot. **(E)** Beta-carotene dynamics for representative intermediate knockouts, colored by the size of growth defect. The knockouts selected were: b1779, b2277, b2779, b0432, b0721, and b3919. The no knockout (NK) curve is shown for reference (green dashed line). **(F)** Heatmaps of peak beta-carotene concentration and rise time. We swept the promoter strength of the rate-limiting enzyme (crtE) across four orders of magnitude. The other enzymes (crtB, crtI, crtY) had fixed promoter strengths equal to those used in the knockout screen.

Pathway simulations (Figure 3B) indeed predict *E. coli* growth in all tested carbon sources, but glucaric acid production was observed only in galactose and glucose, both of which enter glycolysis upstream of glucose-6-phosphate. For all carbon sources except galactose and glucose, the predicted growth rates match those reported previously in the literature^39^, ranging from 0.25 1/hr up to approximately 0.8 1/hr. Growth on galactose and glucose resulted in a 10.3% and 9.3% drop in growth rate, respectively, due to diversion of central carbon flux towards glucaric acid production. These simulations serve as a computational validation of the consistency of our approach and demonstrate its ability to predict dynamics of heterologous pathways in a media-specific fashion.

### E. Impact of gene deletions on pathway dynamics

To test the ability of our approach to predict metabolite dynamics under genetic perturbations, we ran a genome-scale knockout screen on the *E. coli* model, and predicted the dynamics of the beta-carotene pathway (Figure 2C) in each condition. To this end, we zeroed out the flux bounds for the reaction associated to each knockout, and then ran our algorithm with the modified genome-scale model. As in the previous section, this requires re-training the machine learning FBA surrogates for each of the 1,515 knockouts, which in this case involves the generation of large scale data for training, totalling over 757,500 samples computed through off-line FBA simulations.

We found that our knockout simulations predicted the same set of growth essential genes as the wild type iML1515 model, with 12.3% of enzymatic genes being essential with a growth defect above 50% from the wild type (Figure 3C). In line with expectations, beta-carotene production was observed only in the nonessential knockouts. Examination of beta-carotene dynamics across all nonessential knockouts (*N* = 1, 310) suggests that the large majority do not affect the temporal trajectory of product concentration (Figure 3D). However, the simulations also identified a small set of 25 nonessential knockouts associated with intermediate growth defects, ranging from 6% to 18% from the wild type (Figure 3C), that had a pronounced effect on beta-carotene dynamics (Figure 3D). Among these nonessential knockouts, we found a larger growth defects led to a more pronounced peak in beta-carotene concentration (Figure 3E).

Since our approach also allows simulation of pathway dynamics in response to genetic perturbations in the heterologous pathway itself, we selected 6 representative intermediate knockouts from Figure 3C and varied the expression level of the rate limiting enzyme (crtE, shown in Figure 2C) through changes to the strength of its promoter. The results in Figure 3F suggest that peak beta-carotene concentration is primarily sensitive to the promoter strength, while its rise time is primarily affected by the size of the growth defect. Altogether these results underline the utility of our approach to predict metabolite dynamics in response to local and global genetic perturbations.

Despite the computational overhead required to re-train the surrogate models for each of the 1,515 knockouts, we found that our approach provides substantial gains in efficiency as compared to the use of FBA optimization. Even for a relatively high-density training set of *N* = 500 data points, generation of training data took less than 2min/knockout with a negligible time for model training. We performed an additional timing study varying the training set size (Supplementary Table S6). When extrapolated to all 1,515 knockouts, these results suggest that data generation and model training take approximately 49h and 6h, respectively. In contrast, a 24h simulation with the full FBA calculation takes almost 7h/knockout, totalling over 10,000h of computation time for all knockouts.

### F. Host-aware screening of metabolic control circuits

Given that our simulation approach can produce temporal predictions in reasonable computational time, we sought to explore its use as platform for computational screening of metabolic control circuits. Dynamic pathway control has emerged as a powerful tool to build responsive pathways that self-adapt to changes in growth conditions^11,28^. A number of computational approaches have been developed^44,53^ for exploring the space of circuit architectures, but these methods generally overlook the interactions with the host. To illustrate the utility of our approach for host-aware screening of control circuits, we employed two complementary approaches based on parameter sampling for a fixed architecture, and mixed-integer optimization of both parameters and control architectures.

#### a. Large-scale parameter sampling

We first assessed the performance of negative feedback loops from beta-carotene to the promoters producing the pathway enzymes, and compared it to the open-loop case in which all pathway enzymes are constitutively expressed (Figure 4A). The negative feedback acts to downregulate expression of the *crt* enzyme genes in response to beta-carotene, and its has been shown to improve robustness to perturbations^54^.

**FIG. 4.**
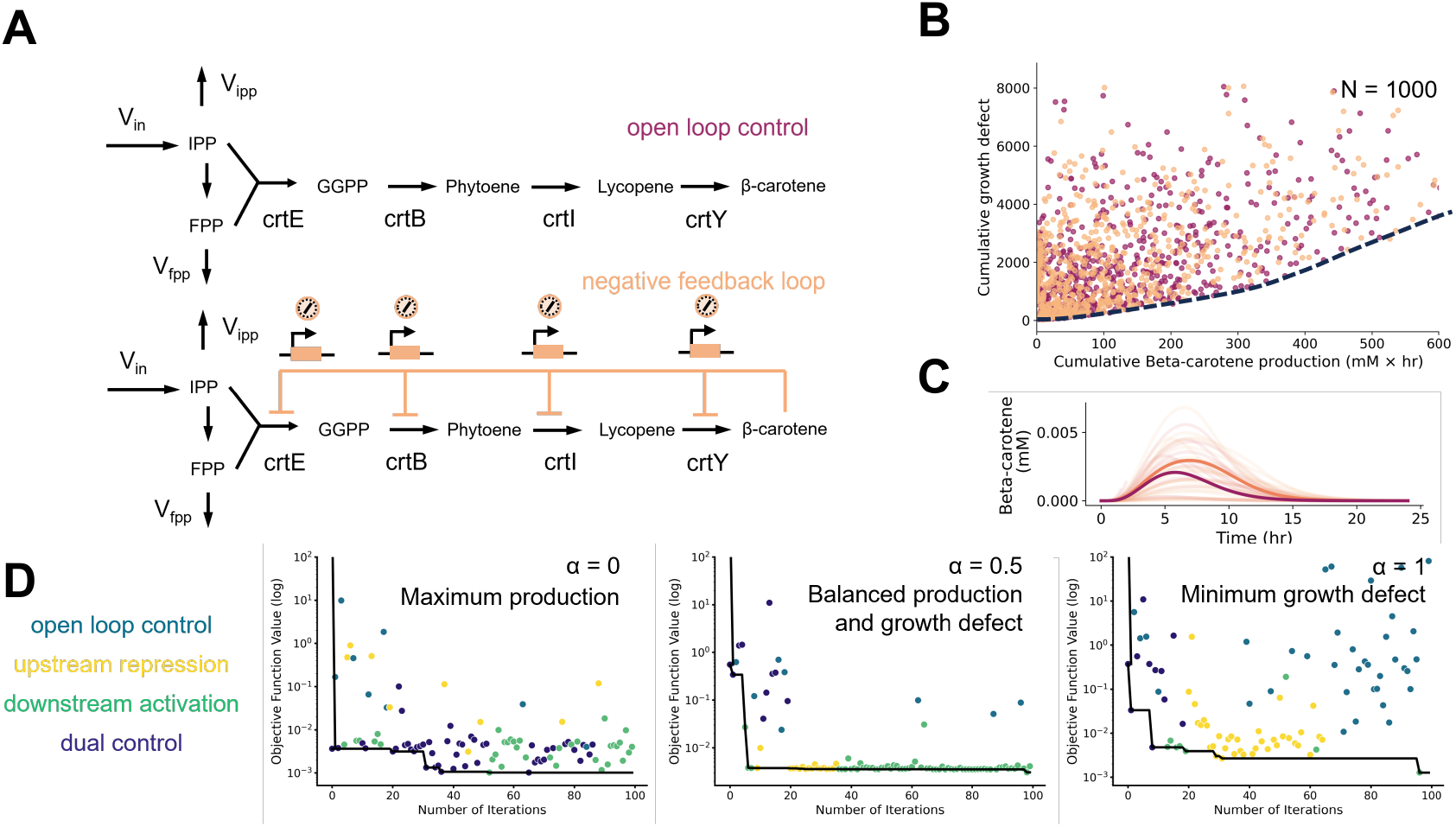
Screening of host-aware control circuits for metabolic pathways. **(A)** Diagram of circuit architectures for beta-carotene pathway in Figure 2C. A negative feedback loop is considered between beta-carotene and the expression of all pathway enzymes via a metabolite-responsive transcription factor. Following the design from Borkowski *et al* ^37^, we considered the four enzymes expressed in an operon with variable ribosomal binding site (RBS) strengths, modelled through four parameters *k*_*i*_ and a single regulatory threshold parameter (*θ*), as shown in Eq. 3 of the main text. **(B)** Random sampling of parameter space. Latin hypercube sampling was used to produce *N* = 1, 000 parameter sets for both open loop control (4-dimensional; no *θ* sampling) and negative feedback (5-dimensional). Bounds for the parameter search psace are given in Supplementary Table S3. Host-pathway simulations were run for 24h and then scored with the cumulative beta-carotene production and cumulative growth defect, defined as 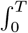 Bcar(*t*) d*t* and 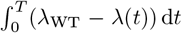, respectively; *λ*_WT_ is the wild type growth rate. An trade-off front emerges between production and growth defect (dashed black line). **(C)** Beta-carotene production dynamics. The median curves are darker; lighter curves are sampled randomly from ±1 standard deviation around the median beta-carotene timecourse. The negative feedback architecture (orange) tends to increase peak beta-carotene concentration when compared to the open loop (red). **(D)** Control circuit screening for glucaric acid production using mixed-integer optimization of both regulatory architectures and parameters. Show are sample runs of a Bayesian optimization loop^41^ applied to the host-pathway model for various weights *α* in Eq. (4) of the main text; the scaling factor was set to *σ* = 1, 256. Points are colored by their respective architectures (shown in Figure 2A) and the black line traces the best architecture found throughout the optimization run. Bounds for the parameter search space are given in Supplementary Table S3.

Following the notation in our general model in Eq. (1), we assumed that all enzymes are expressed at a rate:

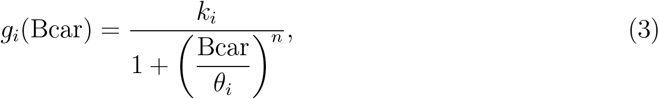

where Bcar is the concentration of beta-carotene, and (*k*_*i*_, *θ*_*i*_) are the promoter strength and regulatory threshold, respectively. The Hill coefficient *n* is set to 2. As noted in Figure 2C, the beta-carotene model results in a transient production and growth defect that returns to the wild type conditions as the precursors are depleted. We thus performed large-scale Latin Hypercube sampling (*N* = 1, 000) of the (*k*_*i*_, *θ*_*i*_) space and scored each design according to the cumulative beta-carotene production and drop in growth rate, both computed as temporal integrals of the simulated dynamics. The results in Figure 4B show the emergence of a trade-off across designs, with high producers resulting in larger drop in growth rate. This trade off is qualitatively similar to the relationship between circuit capacity and growth rate reported by Borkowski et al^37^. We also found that the negative feedback loop increased the median beta-carotene production timecourse (Figure 4C).

#### b. Global optimization of control circuits

As a final test of our approach, we used it to automatically search for optimal control parameters and architectures, using the glucaric acid pathway as a test case. We restricted the search to three genetic feedback architectures in addition to open loop control (Figure 4D) and employed a recent Bayesian Optimization algorithm to search over the continuous-discrete design space efficiently^41^. This is a challenging optimization problem that requires simulating the pathway dynamics at many points in the input space, and thus a ideal test to assess if our simulation approach could be deployed within such computationally intensive tasks.

Since the glucaric acid pathway leads to a permanent growth defect (Figure 2B), we constructed an objective function *J* that weighs the drop in steady state growth rate and the cumulative production of glucaric acid:

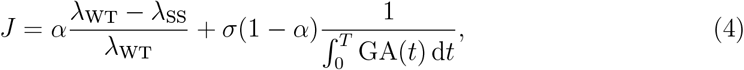

where *λ*_SS_ and *λ*_WT_ are the steady state growth rate of the production and wild type strain, respectively, and GA is the concentration of glucaric acid. The parameter *σ* is scaling factor to ensure both terms are in the same magnitude, and *α* is a weight to control the balance between growth defect and production. The optimization results in Figure 4D show that the optimizer can effectively single out optimal design; for a fixed value of *α*, each optimization run (*N* = 100 iterations, 10h of simulation time per iteration) took 4.1h of computations. In all considered cases we observed the optimizer converging to an optimal control circuit.

## III. DISCUSSION

The interplay between engineered pathways and host metabolism is a key determinant of production phenotypes. Here, we developed a strategy to predict the dynamics of heterologous pathways coupled with the metabolism of the host. Our approach bridges an important gap in the literature by reconciling kinetic and genome-scale models, two well-adopted metabolic modelling paradigms, into a computational tool to probe the impact of environmental and genetic perturbations on the dynamic cross-talk between pathway and host. Our approach can be readily employed to simulate heterologous pathways coupled with an off-the-shelf GEM, without the need to extend it with the pathway of interest, or manual calibration of the biomass objective function. The methodology can also readily accommodate alternative objective functions that trade off biomass against production or other cellular objectives, as well as optimization strategies designed for mutant strains that do not satisfy the same optimality principle as the wild-type, such as MOMA^55^ or ROOM^56^. The approach can be employed to explore the impact of pathway dynamics on production phenotypes, for example by studying pathway bottlenecks due to metabolite accumulation, or the onset of toxicity or stress responses upon enzyme overexpression. This may also provide insights on dynamic control strategies that can potentially mitigate the impact host-pathway interactions and enable the identification of control architectures with improved performance^28^. Such control strategies have been shown to help manage the cross-talk between gene circuits and their host^57,58^, and they may also provide benefits in pathway engineering.

Moreover, our method can serve as a tool for genome-scale knockout screens to identify genomes with improved production phenotypes. While host-pathway interactions are normally seen as deleterious for production, it is well known that genetic knockouts can cause unexpected effects on cell physiology; for example, a large-scale study showed that a small fraction of double yeast knockouts can improve fitness^59^, while multiplexed CRISPR screens have found combinations of knockouts with improved metabolic phenotypes^60,61^. Such findings pave the way for the use of such synergistic genetic interactions as a novel strategy to improve production. Our simulation approach could be thus employed in combination with well-adopted strain optimization tools such as OptKnock or OptForce^62,63^ to identify genetic edits with improved pathway dynamics.

Another promising application of this work is the study of metabolic burden^64^. The synthetic biology literature has so far focused mostly on expression burden and its impact on cell physiology^15,16,65,66^. In a seminal work, Borkowski and colleagues^37^ employed a cell free platform to show that metabolic burden, as quantified by a decreased pathway titer, could not be explained solely by the increased competition for gene expression resources. This hints to a metabolic source of burden that is challenging to define and quantify, which is an area of research where our approach can help drawing new hypothesis on the sources and impact of metabolic burden.

A key enabler in our approach was the use of machine learning models to replace costly FBA calculations. This led to substantial gains in simulation speed that would have otherwise made the FBA-ODE integration too slow for practical use. However, there is additional room for further improvements to computational efficiency. The main bottleneck is the generation of data for training the surrogate machine learning models, which must be instantiated separately for every modification of the GEM, including knockouts and changes to media conditions. A transfer learning approach that selects minimal training data computed on modified GEMs could potentially speed up this process^67^. Previous work has used active learning approaches to train surrogate models with fewer expensive computational simulations^68^ or used iterative parallel computing across multiple CPU cores to speed up the training data generation^69^.

Another drawback is that model integration must be done in a pathway-specific fashion. We have illustrated how to define the boundary fluxes and modify the kinetic model so that the FBA steady state constraints are respected. In general, however, such model modifications are pathway-dependent and thus may require substantial user input. In the Supplementary Table S7 we have enumerated some representative pathway topologies to ease the application of our strategy across other pathways with more complex branch point topologies than the considered case studies.

Our work contributes to the emerging area of hybrid metabolic modelling that attempts to expand the predictive power of traditional approaches and incorporate a broader range of cellular processes relevant for pathway engineering. Recent examples include the use of machine learning to augment the predictive power of GEMs^70,71^, integration of FBA and bioprocess models^22,23^, machine learning for improved genome-scale kinetic modelling^72^, and expanding the boundaries of GEMs to include signalling pathways^73^. We anticipate this area will continue to grow and draw inspiration from machine learning techniques as practitioners strive to balance mechanistic and data-driven approaches to improve production across a range of pathway engineering tasks^74^.

## IV. METHODS

### A. Simulation algorithm

We can express the our simulation loop in pseudocode as:

#### Algorithm 1

FBA-ODE simulation loop

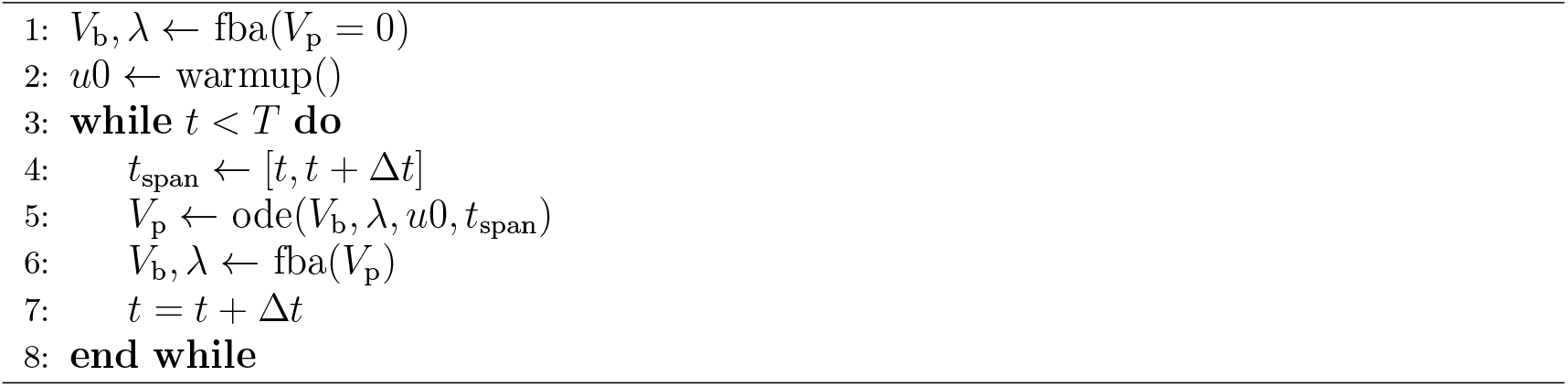

The loop begins by initializing the boundary flux vector *V*_b_ and the growth rate *λ* using FBA with the pathway flux *V*_p_ set to zero. We obtain the intial condition concentration values from the ODE using the warmup routine (for details, see Section IV C). We consider both models operating in distinct alternating timesteps and divide the simulation interval [0, *T*] into *N* equispaced subintervals of length Δ*t*. Within each subinterval [*t, t* + Δ*t*] we solve the ODE model for the heterologous pathway:

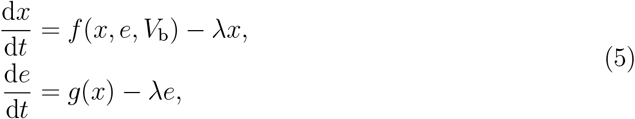

where the function *f* (*x, e, V*_b_) depends explicitly on the boundary fluxes (*V*_b_) feeding into the pathway from the host. The enzyme and substrate concentration values at the end of the integration time *t* + Δ*t* are used to compute the pathway flux *V*_p_. The pathway flux *V*_p_ is passed to the FBA solver as a constraint on the added sink reaction 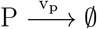:

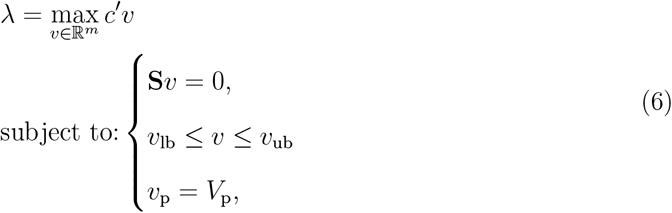

The updated FBA problem re-computes the optimal flux vector and growth rate *λ* to start a new integration cycle of the ODE model. The time *t* is updated every loop iteration to move it forward one interval with length Δ*t*. In all our simulations we employed Δ*t* = 1s as a tradeoff between coarse-graining the FBA optimization and computational efficiency.

### B. Models considered in case studies

All models were implemented in the Julia scientific programming language, using the COBREXA.jl package for GEM analysis^75^ and DifferentialEquations.jl for fast ODE integration^76^.

#### 1. Genome-scale model for the host

In both case studies we employed the iML1515 model for *Escherichia coli* ^39^. An additional reaction was added to the model to represent the dynamic pathway with stoichiometry of -1.0 for all precursors and no product. For the glucaric acid pathway, we employed the default growth media conditions in the iML1515 model (glucose as carbon source with a maximal uptake rate of 10mM/h). For the beta-carotene pathway, we found an experimentally determined wild type growth rate of 0.65 1/h in the literature^37^. We adjusted the medium conditions in the FBA to a limited fructose source by setting a constraint on the fructose influx (−7.5 mM/h) that resulted in an equivalent growth rate when the heterologous pathway was not induced.

#### 2. Glucaric acid pathway

The ODE model for the glucaric acid pathway in Figure 2A was taken from literature^44^ and modified to allow a variable influx and growth rate passed from FBA. The model includes three metabolites (glucose-6-phosphate, fructose-6-phosphate, and myoinositol) and two enzymes (Ino1 and MIOX). The other steps in the conversion of myoinositol to glucaric acid were assumed to be non-rate-limiting and were not included in the model. All kinetic parameters were taken from the BRENDA database^77^. The model equations and parameter values are included in Section S1 of the Supplementary Information.

The boundary flux (*V*_in_) in this case is defined as the sum of 11 reactions that produce or consume glucose-6-phosphate: TRE6PH, PGMT, HEX1, AB6PGH, TRE6PS, FRULYSDG, GLCptspp, G6PP, G6Pt62pp, and BGLA1. To model the consumption of G6P by the glucaric acid pathway, a new forward reaction was added to the iML1515 model (*V*_p_, which consumes 1 mol of G6P, has no products, and is constrained to run forward to prevent flux being drawn backwards through the production pathway.

#### 3. Beta-carotene pathway

For the beta-carotene pathway in Figure 2C, we constructed a kinetic model including the four Crt enzymes and six metabolites (FPP, IPP, GGP, phytoene, lycopene, and beta-carotene). We assumed Michaelis-Menten kinetics for all enzymes with kinetic parameters taken from BRENDA. For those *k*_cat_ values that were not available, we employed a deep learning predictor^52^. The model equations and parameter values are included in Section S2 of the Supplementary Information.

Since the pathway has two precursors (farnesyl pyrophposhate, FPP, and isopentyl pyrophosphate, IPP), the boundary flux vector *V*_b_ has several components. We define a boundary influx into IPP (*V*_in_) as the sum of the reactions IPDDI and IPDPS. We define an efflux to IPP (*V*_ipp_) as the sum of the reactions UDCPDPS, OCTDPS, DMATT, IPDPS with appropriate stoichiometric weighting. FPP is entirely produced by IPP and its efflux (*V*_fpp_) is defined by the sum of the reactions UDCPDPS, HEMEOS, and OCTDPS. A new forward reaction (*V*_p_) was added to the iML1515 model to represent consumption of FPP and IPP by the beta-carotene pathway. This reaction consumes 1 mol of FPP and 1 mol of IPP, has no products, and is constrained to run forward to prevent flux being drawn backwards through the production pathway.

### C. Considerations for model integration

#### Balancing fluxes at the model boundary

Since the precursor and dynamic pathway flux *V*_p_ are included in both models, the ODE must satisfy the steady-state constraint imposed by FBA at each iteration of the simulation loop; that is, the reaction fluxes that produce and consume the precursor(s) must balance out. The balancing equations are pathway-dependent and described for each of our case studies next.

The glucaric acid pathway required no modifications to the ODEs as it is the simplest case in which one boundary flux feeds into the precursor G6P and one pathway flux consumes it. The flux balance is therefore simply:

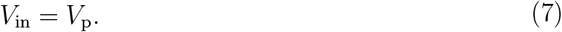

The precursor equation remains unchanged from the literature model:

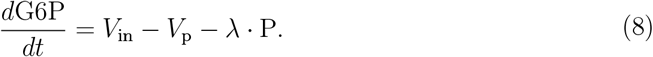

For the beta-carotene pathway, both precursors (FPP and IPP) combine in an equimolar reaction to produce the pathway flux *V*_p_. We define three components of the boundary flux vector: *V*_in_ and the two precursor effluxes *V*_fpp_ and *V*_ipp_. The unbalanced precursor equations are as follows:

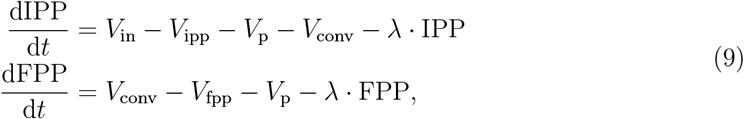

where *V*_conv_ is the flux converting IPP to FPP, which is included in the FBA but is not a boundary flux and therefore not passed to the ODE. There is no inherent constraint on the ODE which ensures that the boundary fluxes and *V*_conv_ must satisfy the steady-state assumption, so we must impose one by expressing *V*_conv_ in terms of the boundary fluxes. Taking a flux balance of each precursor, we obtain the following equations:

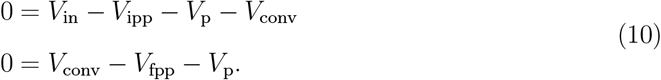

We rearrange the second equation to find an expression for *V*_conv_ in terms of only the boundary and pathway fluxes:

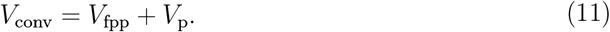

We substitute this expression into the the precursor equations in Eq. (9) and obtain:

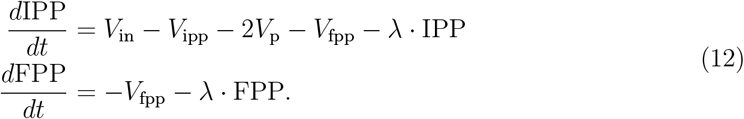

This modification to the ODE model ensures that the boundary fluxes (*V*_in_, *V*_fpp_, and *V*_ipp_) satisfy the steady-state constraint at each iteration of the simulator loop. These ODE modifications are pathway-specific and in Supplementary Table S7 we show the balancing equations for several common topologies encountered in applications.

#### Warm-up routine for initial conditions

To ensure that the initial conditions of the ODE lead a to feasible flux *V*_p_ for the FBA problem, we created a warm-up routine which iteratively tries various initial conditions using a Bayesian optimization method^78^. These initial conditions are then run for 500 iterations of the simulator (500 seconds) and each iteration is checked for FBA feasibility. If initial conditions are not valid for all 500 iterations, the objective function of the Bayesian optimizer is heavily penalized:

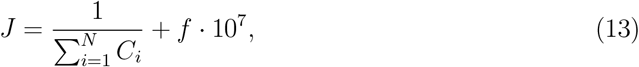

where *f* is the binary feasibility flag (1 if infeasible) and *C*_*i*_ are the concentrations of native metabolites, i.e. G6P and F6P for the glucaric acid pathway, and FPP and IPP for the beta-carotene pathway. We assume that the initial concentrations of heterologous enzymes and pathway metabolites are nil, as the host does not produce them natively before the pathway is induced. The warmup routine runs for 1000 initial condition values and the initial conditions which achieve the lowest value of *J* are selected for further simulation. If no feasible conditions are found, i.e. when the minimum objective function value is greater than 10^7^, no further simulation runs. For the Bayesian optimization loop, we employed a Tree of Parzen Estimator (TPE) algorithm implemented in the TreeParzen.jl Julia package. Expected Improvement was used as the acquisition function and initial conditions for the native metabolites (G6P, F6P, FPP, and IPP) were drawn from uniform priors with a maximum of 0.75mM (glucaric acid) and 10mM (beta-carotene), set by integration the ODE to steady state without the pathway induced, i.e. without heterologous enzymes.

### D. Machine learning surrogate models

To produce simulations in feasible computational time, we substituted all FBA-related steps with pre-trained machine learning surrogates. We split the surrogates into three supervised tasks: a binary classification problem to predict whether FBA is feasible for a given pathway flux (*V*_p_), a linear regression between pathway flux (*V*_p_) and optimal growth rate (*λ*), and a nonlinear regression to predict the boundary fluxes (*V*_b_) from the pathway flux (*V*_p_).

Once trained, we can implement the machine learning surrogates in the existing algorithm loop as follows:

#### Algorithm 2

Surrogate-ODE Optimization Loop

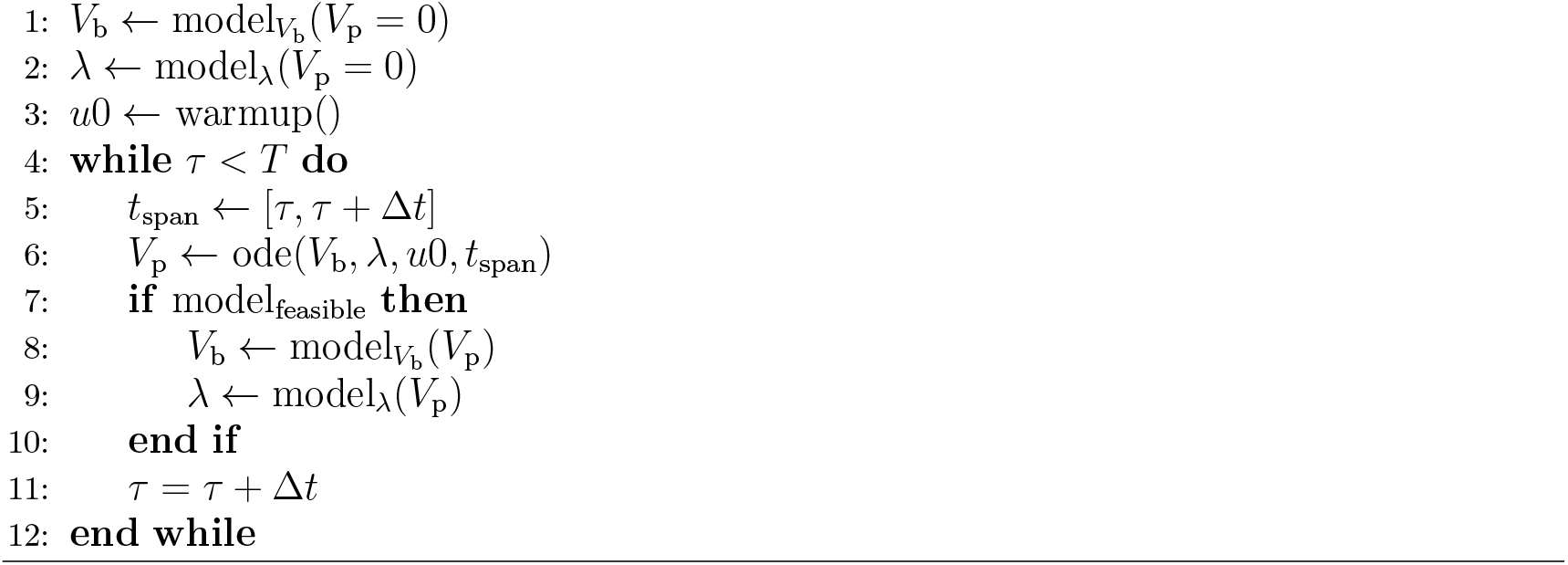

Instead of computing *λ* and the boundary flux directly using FBA, we first check if FBA finds a particular *V*_p_ feasible and then generate the *V*_b_ and *λ* values from their respective models. Neural networks were trained using the Flux.jl package and linear models using the GLM package. The exact training data generation routines and model specifications for each case study are given next.

#### Glucaric acid model

We generated training data by sweeping the pathway flux (*V*_p_) from 0 to 10 mM/h and sampled 500 linearly spaced values to constrain the FBA problem. Generation of the training data and model training took around 5 minutes on a standard laptop. We randomly split the data into training and testing using a 80:20 split, and trained a logistic regressor to predict model feasibility. We consistently achieved test accuracy above 95% (Supplementary Figure S1A). To regress the optimal growth rate and boundary fluxes, we first we filtered the training set to include only feasible data points, and then trained a linear regressor that achieved *R*^2^ of 0.99 or above (Supplementary Figure S1B). To regress the boundary fluxes, we trained a feedforward neural network with 3 layers, 500 units per layer, and ReLu activation function for 1500 epochs (Supplementary Figure S1C). When comparing the predicted results to ground truth, we find a good fit that avoids overfitting to numerical noise (Supplementary Figure S1D). We trained *N* = 5 training repeats and achieved consistently high performance (Supplementary Table S5). We also confirmed that model performance degraded with smaller training set sizes (Supplementary Figure S3). We incorporated the machine learning models into the loop as a replacement for FBA, which improved the model run time by several orders of magnitude. For example, model runs that took >1000 seconds in the standard loop were accelerated to 12 seconds with the surrogate models (see Supplementary Figure S2). Models and training data objects were serialized to .jls files to minimize retraining time. For the variable growth conditions in Figure 3A–B, we modified the uptake reactions for all carbon sources to limit influx to a single source and regenerated training data for each carbon source. All models were retrained for a total of 15 conditions (*N* = 45 new models).

#### Beta-carotene model

Training data was generated by varying the pathway flux (*V*_p_) from 0 to 1mM/h and sampling 500 linearly spaced values to constrain the FBA problem. Generation of the training data set and model training took around 5 minutes on a standard laptop. We held out 20% of the training set for model testing. We trained a logistic regression model to predict model feasibility. We trained a linear regression model to predict growth rate. We found a linear relationship between *V*_p_ and all boundary fluxes, so to speed up training time we trained three linear regression models to predict the boundary flux components *V*_in_, *V*_ipp_, and *V*_fpp_. All linear models obtained an *R*^2^ of 0.99 or better and the logistic regression obtained a test accuracy of 0.99. Models and training data objects were serialized to .jls files to minimize retraining time. For the genome-wide knockout screen in Figure 1C–E, the FBA model was modified by knocking out each gene and generating new training data in each case, and then retraining all models (*N* = 1, 515 knockouts, *N* = 7, 575 models).

## ACKNOWLEDGEMENTS

CM and DAO were supported by the United Kingdom Research and Innovation (grant EP/S02431X/1, UKRI Centre for Doctoral Training in Biomedical AI).

## Supplementary Information

### S1. KINETIC MODEL FOR THE GLUCARIC ACID PATHWAY

The model for the glucaric acid pathway in Figure 2A was adapted from previous work^41^.

The mass balance equations are:

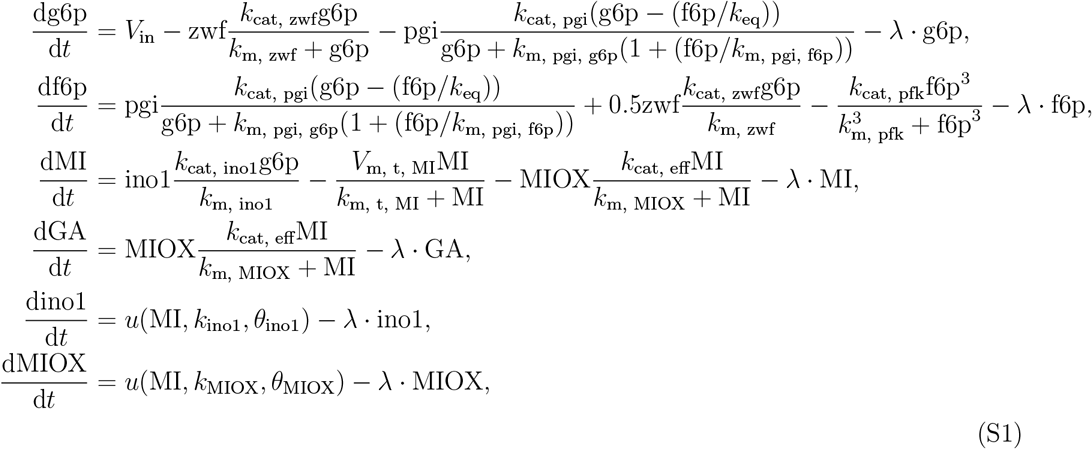

where *V*_in_ is a constant influx to the pathway, Ino1 and MIOX are the heterologous enzymes, and g6p, f6p, MI (myoinositol), and GA (glucaric acid) are the pathway intermediates. The enzyme parameters *k*_cat_, *k*_eq_, *k*_m, x_ and *k*_m, y_ are all fixed kinetic parameters specific to each enzyme. The effective substrate activation constant *k*_cat, eff_ has two additional activation kinetic parameters *k*_a_ and *a* which must be specified:

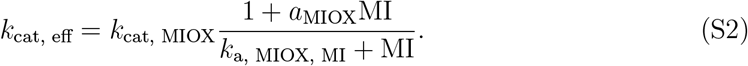

The influx *V*_in_ and cell growth rate *λ* are input from the FBA model. The function *g*(*x, k, θ*) describes the genetic control topology at the enzyme’s promoter. There are three possible topologies at each promoter, open loop control, activation, and repression:

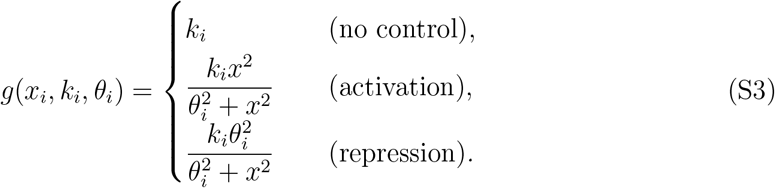

The possible architectures are limited to three architectures which include only negative feedback loops, in addition to an open-loop control with no dynamic feedback loops (see Figure 1A). This architecture limitation was made to remove the possibility of undesirable multistability or oscillatory behavior. All parameter values are given in Table S1.

### S2. KINETIC MODEL FOR THE BETA-CAROTENE PATHWAY

We built a model for the beta-carotene pathway in Figure 2C. We modeled the four enzymes in the pathway (crtE, crtB, crtI, crtY) along with six metabolites: FFP, IPP, GGP, phytoene, lycopene, and beta-carotene. A constant influx to the pathway *V*_in_ and a growth rate *λ* are set as parameters in the ODE. The first reaction in the pathway follows Michaelis-Menten kinetics with two substrates:

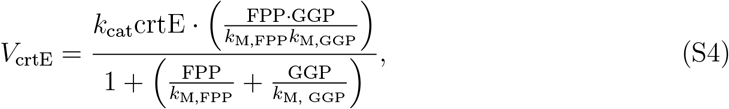

where *k*_M,FPP_ and *k*_M,GGP_ are substrate-specific enzyme affinity parameters. All other enzymes are assumed to follow standard Michaelis-Menten kinetics:

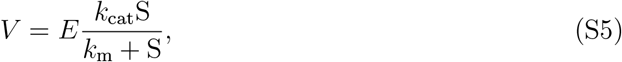

where *E* is the enzyme concentration, *S* is the substrate concentration, and *k*_cat_ is the turnover number. We sourced *k*_m_ values and averaged available *k*_cat_ numbers from BRENDA^77^; however, the turnover numbers were not available for some enzymes and often available only for unrelated organisms. We instead used a deep learning model trained on all available enzyme kinetic information to predict *k*_cat_ values from enzyme amino acid sequence^52^. For those *k*_cat_ values available in BRENDA, the deep learning model delivered predictions in the same order of magnitude.

The substrate equations can thus be written as:

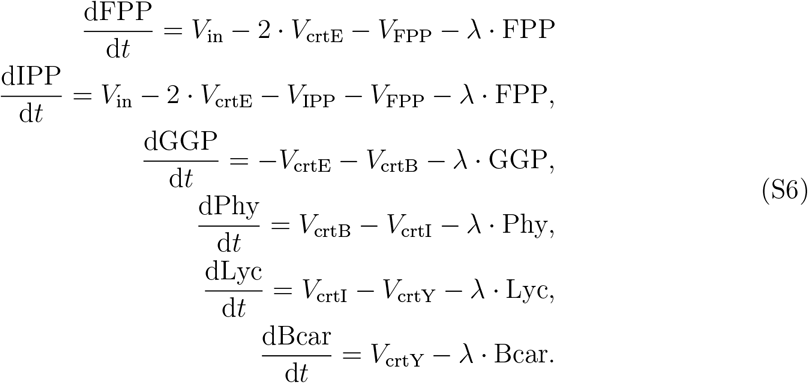

The equations for all four enzymes follow standard open-loop production with a promoter strength parameter *k*_*E*_. For example, for the first enzyme in the pathway, the equation is

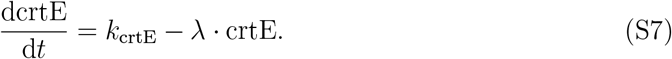

Table S2 contains all kinetic parameter values for the ODE model.

### SUPPLEMENTARY TABLES

**Supplementary Table S1.** Kinetic parameters of the glucaric acid pathway model. The parameters *λ* and *V*_in_ were used only in the warm-up routine to find initial conditions; in all other cases, both parameters were iteratively computed through the FBA-ODE loop.

| Parameter | Value | Units | Parameter | Value | Units |
| --- | --- | --- | --- | --- | --- |
| $\lambda$ | 2.77E-5 | 1/s | $n_{\text{Pfk}}$ | 3 | N/A |
| $V_{\text{in}}$ | 0.1656 | mM/s | $k_{\text{cat, ino1}}$ | 0.2616 | mM/s |
| $k_{\text{cat, pgi}}$ | 0.8751 | mM/s | $k_{\text{m, ino1, g6p}}$ | 1.18 | mM |
| $k_{\text{eq, pgi}}$ | 0.3 | mM | $V_{\text{m, t, MI}}$ | 0.045 | mM/s |
| $k_{\text{m, pgi, g6p}}$ | 0.28 | mM | $k_{\text{m, t, MI}}$ | 15 | mM |
| $k_{\text{m, pgi, f6p}}$ | 0.147 | mM | $k_{\text{cat, MIOX}}$ | 0.2201 | mM/s |
| $k_{\text{cat, zwf}}$ | 0.0853 | mM/s | $k_{\text{m, MIOX}}$ | 24.7 | mM |
| $k_{\text{m, zwf, g6p}}$ | 0.1 | mM | $a_{\text{MIOX}}$ | 5.422 | N/A |
| $k_{\text{cat, pfk}}$ | 2.615 | mM/s | $k_{\text{a, MIOX, MI}}$ | 20 | mM |
| $k_{\text{m, pfk, f6p}}$ | 0.16 | mM | | | |

**Supplementary Table S2.**
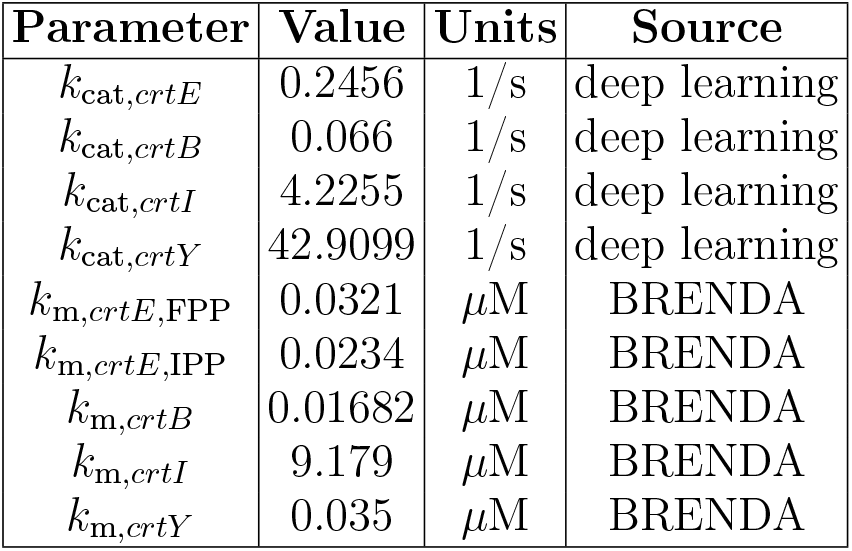
Kinetic parameters of the beta-carotene pathway model. For those parameters absent from the BRENDA database, we employed the deep learning predictor from Li *et al* ^52^.

**Supplementary Table S3.** Parameter values and bounds for simulations in Figures 2–4 in the main text.

| Figure | Model | Simulation | Parameter Values |
| --- | --- | --- | --- |
| 2 | glucaric acid | sample simulation | dual control architecture<br>$k_1 = 2 \times 10^{-5}$<br>$k_2 = 2.2 \times 10^{-3}$<br>$\theta_1 = 3.3$<br>$\theta_2 = 1.0$ |
| 2 | beta-carotene | sample simulation | $k_1 = 6.5 \times 10^{-6}$<br>$k_2 = 2.3 \times 10^{-2}$<br>$k_3 = 3.2 \times 10^{-2}$<br>$k_4 = 1.2 \times 10^{-2}$ |
| 3 | glucaric acid | medium conditions | open loop architecture<br>$k_1 = 4.1 \times 10^{-5}$<br>$k_2 = 2.27 \times 10^{-4}$ |
| 3 | beta-carotene | knockouts | $k_1 = 2.41 \times 10^{-7}$<br>$k_2 = 9.7 \times 10^{-5}$<br>$k_3 = 9.8 \times 10^{-5}$<br>$k_4 = 3.67 \times 10^{-4}$ |
| 4 | beta-carotene | large-scale parameter sampling | $10^{-8} \leq k_1 \leq 10^{-6}$<br>$10^{-7} \leq k_2 \leq 10^{-5}$<br>$10^{-6} \leq k_3 \leq 10^{-5}$<br>$10^{-5} \leq k_4 \leq 10^{-4}$<br>$10^{-5} \leq \theta \leq 10^1$ |
| 4 | glucaric acid | global optimization of control circuits | 4 architectures:<br>open loop control<br>upstream repression<br>downstream activation<br>dual control<br>$10^{-7} \leq k_1 \leq 10^{-3}$<br>$10^{-7} \leq k_2 \leq 10^{-3}$<br>$10^{-7} \leq \theta_1 \leq 10^1$<br>$10^{-7} \leq \theta_1 \leq 10^1$ |

**Supplementary Table S4.**
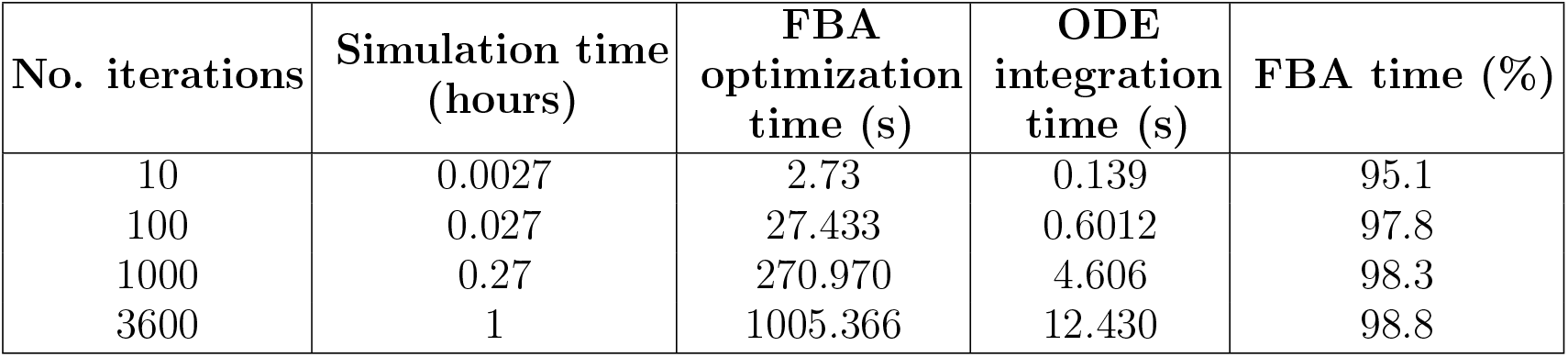
Runtime analysis of integrated FBA and ODE simulations. Study was done on the glucaric acid pathway using an iteration timestep of 1s to balance computational costs and coarse-graining the FBA predictions.. The simulation was run following Algorithm 1 in the Methods. Details can be found in code repository (see Code Availability section).

**Supplementary Table S5.** Performance of machine learning models. The machine learning models for the glucaric acid model were run *N* =5 times with different random initializations and the accuracy metrics for each model reported with standard deviation across repeats.

| Model | Test Accuracy | Model $R^2$ | Training MSE | Test MSE |
| --- | --- | --- | --- | --- |
| Feasibility | $99.8 \pm 0.005$ | N/A | N/A | N/A |
| Growth Rate | N/A | $0.999 \pm 1.62\text{E-}7$ | $3.45\text{E-}33 \pm 3.86\text{E-}33$ | $3.56\text{E-}9 \pm 4.88\text{E-}9$ |
| Pathway Influx | N/A | N/A | $0.0124 \pm 0.0198$ | $0.0167 \pm 0.0226$ |

**Supplementary Table S6.** Computational cost for generation of training data. We conducted a timing study was conducted on the glucaric acid pathway with glucose as carbon source. Fresh training data for growth in glucose was generated *N* =5 times and the three machine learning models were re-trained from scratch *N* = 5 times to compute average times and their standard deviation.

| Training set size | Training data generation time (s) | ML model training time (s) |
| --- | --- | --- |
| 500 | $117.2 \pm 8.60$ | $14.27 \pm 3.87$ |
| 100 | $27.9 \pm 3.24$ | $4.12 \pm 0.46$ |
| 50 | $21.1 \pm 2.85$ | $3.17 \pm 0.29$ |
| 25 | $8.57 \pm 0.45$ | $2.36 \pm 0.095$ |

**Supplementary Table S7.**
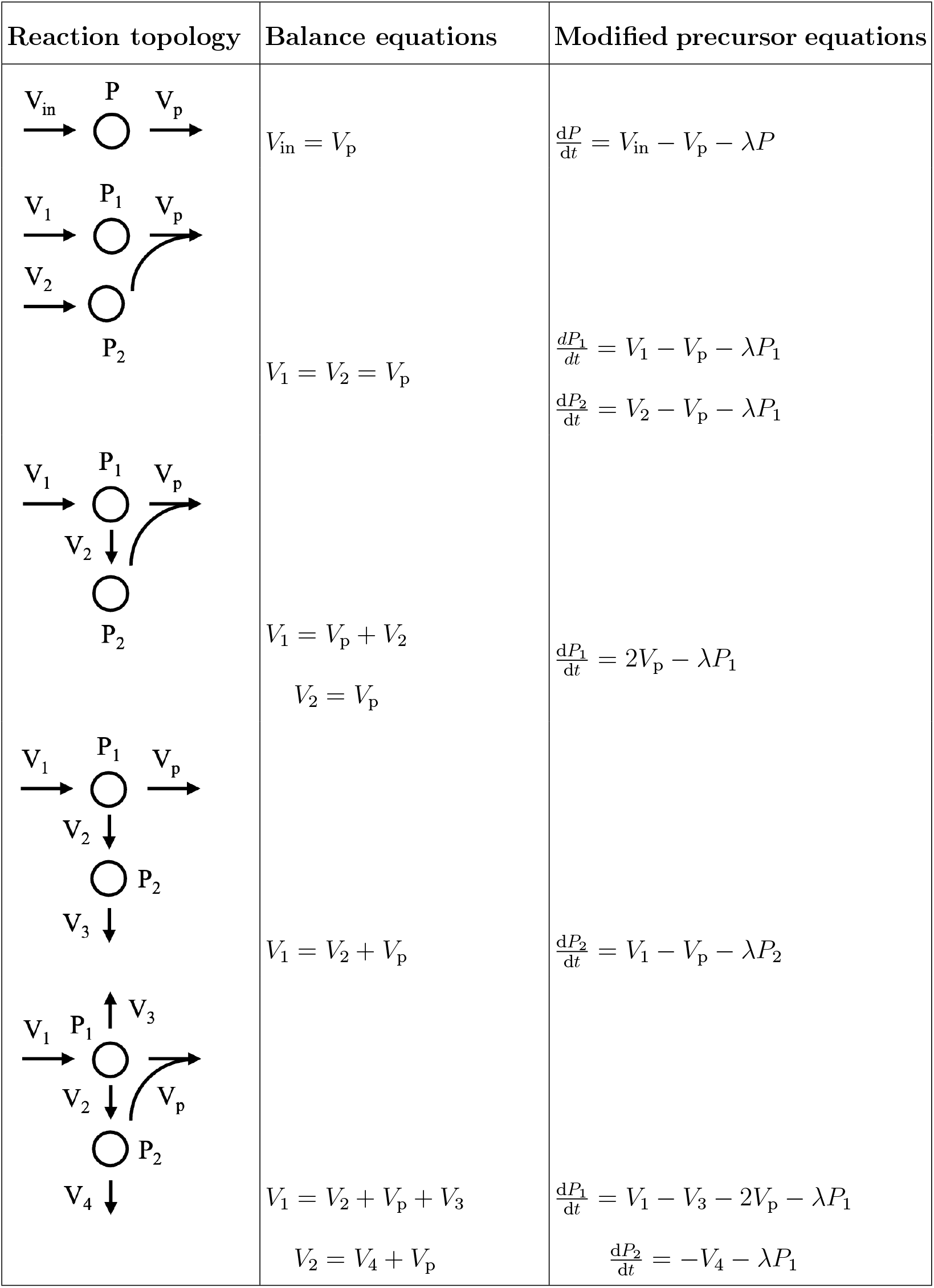
Reaction topologies and their associated balancing equations. Balancing the pathway fluxes at the boundary for various reaction topologies requires algebraic substitutions to modify the ODE model so that they maintain the steady state assumption at each timestep. Table shows only the precursor equations that need to be modified. The glucaric acid and beta-carotene pathways in Figure 2 correspond to the first and last topology, respectively.

| Reaction topology | Balance equations | Modified precursor equations |
| --- | --- | --- |
| | $V_{in} = V_p$ | $\frac{dP}{dt} = V_{in} - V_p - \lambda P$ |
| | $V_1 = V_2 = V_p$ | $\frac{dP_1}{dt} = V_1 - V_p - \lambda P_1$<br>$\frac{dP_2}{dt} = V_2 - V_p - \lambda P_1$ |
| | $V_1 = V_p + V_2$<br>$V_2 = V_p$ | $\frac{dP_1}{dt} = 2V_p - \lambda P_1$ |
| | $V_1 = V_2 + V_p$ | $\frac{dP_2}{dt} = V_1 - V_p - \lambda P_2$ |
| | $V_1 = V_2 + V_p + V_3$<br>$V_2 = V_4 + V_p$ | $\frac{dP_1}{dt} = V_1 - V_3 - 2V_p - \lambda P_1$<br>$\frac{dP_2}{dt} = -V_4 - \lambda P_1$ |

### SUPPLEMENTARY FIGURES

**Supplementary Figure S1.**
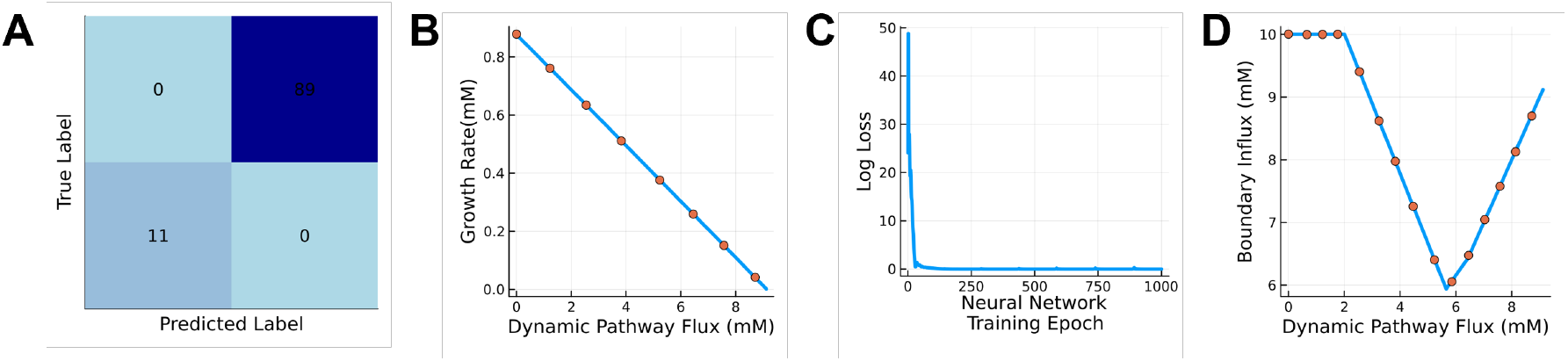
ML surrogate model training. All results in the following parts are based on a model trained on the wild type iML1515 model grown on glucose at default uptake rates of 10mM/h with the boundary flux computed as the *V*_in_ from the glucaric acid model. The dynamic pathway flux *V*_p_ was swept from 0 to 10mM/h and FBA was run to determine feasibility. If feasible, we also computed the growth rate and boundary flux. We split the data generated from FBA (*N* = 500 samples) into a training (80%) and test (20%) set and reserved the test set for model evaluation. **(A)** A logistic regression model trained on a binary feasibility class (1 = feasible, 0 = infeasible) predicts FBA feasibility. FBA is infeasible when no positive growth rate exists within the constrained flux cone. We generated a confusion matrix for a 20% held test set (*N* = 100). The predicted labels were identical to the ground truth labels with zero false positives or false negatives. **(B)** Feasible training samples were selected for further model training. A linear regression model was trained to predict the growth rate from the dynamic pathway flux *V*_p_. The blue line is the ground truth and the orange circles are selected trained model predictions from the held-out test set. **(C)** Training loss for boundary flux neural network. A feedforward neural network with 3 hidden layers, 500 units per layer, and ReLu activation function was trained for 1000 epochs. The loss converged after approximately 250 epochs.**(D)** Boundary flux predictions. The blue line is the ground truth and the orange circles are a subset of test set predictions. The shape of the ground truth curve varies based on the medium conditions and thus cannot be assumed to be linear like the growth rate relationship.

**Supplementary Figure S2.**
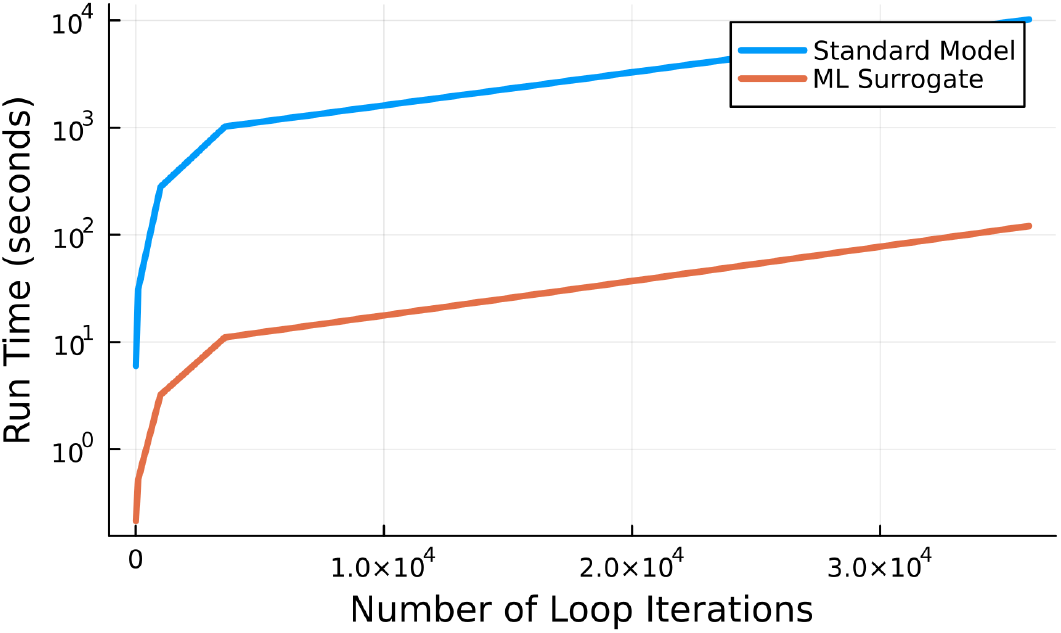
Timing comparison between FBA and ML surrogate. Loop runtime for standard FBA and ML surrogate loop, in seconds (log scaled). Both increase linearly with the number of loop iterations; however, standard FBA becomes rapidly computationally infeasible due to the cost of linear FBA optimization. For comparison, most simulations run for the results require 8.6 × 10^4^ loop iterations and would take > 6.6hrs to run with a standard FBA model, compared to around 5 minutes with the ML surrogate.

**Supplementary Figure S3.**
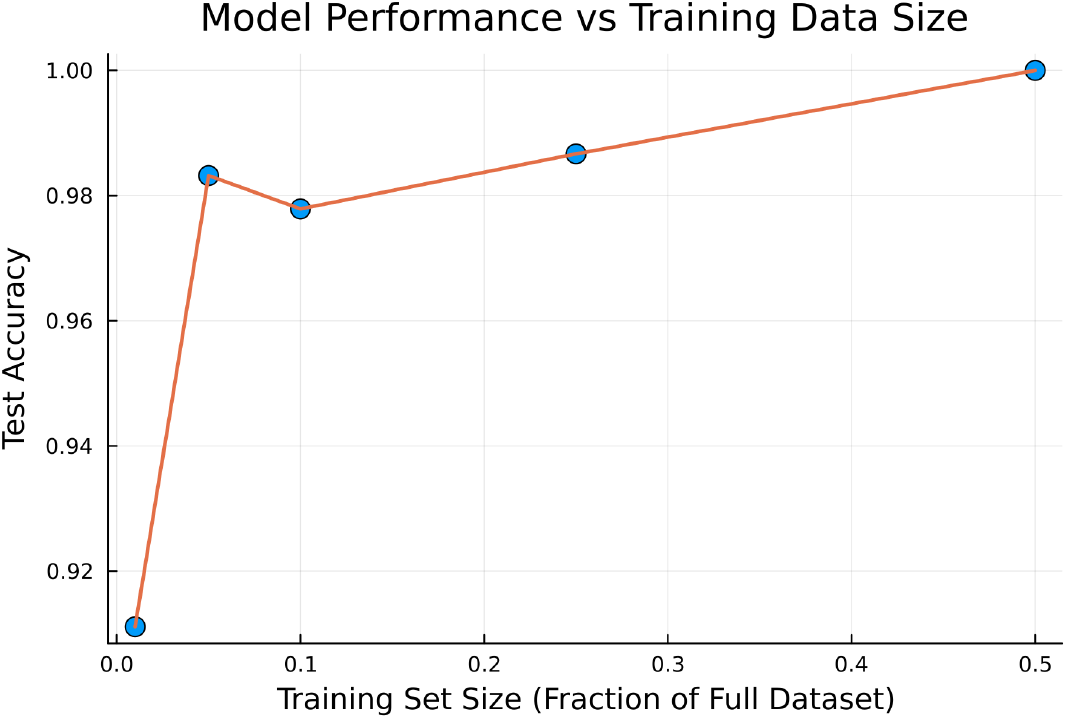
Degradation of feasibility model performance. We reduced the training set size for the feasibility assessment by 50-99% and plotted the test accuracy of the feasibility binary classification. The performance of the regression models also degrades when assessing test MSE but less substantially, likely due to the simple functional form of the prediction task.

